# Selective axonal translation of prenylated Cdc42 mRNA isoform supports axon growth

**DOI:** 10.1101/366369

**Authors:** Seung Joon Lee, Amar N. Kar, Matthew D. Zdradzinski, Priyanka Patel, Pabitra K. Sahoo, Riki Kawaguchi, Byron J. Aguilar, Kelsey D. Lantz, Caylee R. McCain, Giovanni Coppola, Qun Lu, Jeffery L. Twiss

**Affiliations:** Department of Biological Sciences, University of South Carolina, Columbia, SC 29208 USA; Department of Psychiatry, Semel Institute for Neuroscience and Human Behavior, Los Angeles, CA 90095-1761, USA; Department of Anatomy and Cell Biology, Brody School of Medicine, East Carolina University, Greenville, NC 27834 USA; Department of Neurology, Semel Institute for Neuroscience and Human Behavior, Los Angeles, CA 90095-1761, USA

**Author notes:** Corresponding Author: Jeffery L. Twiss, M.D., Ph.D., Dept. Biological Sciences, University of South Carolina, 715 Sumter St., CLS 401, Columbia, SC 29208 USA, email.

**Keywords:** alternative splicing, axon regeneration, growth cone, mRNA transport, geranylgeranylation

## Abstract

The small Rho-family GTPase Cdc42 has long been known to have a role in cell motility and axon growth. The eukaryotic *CDC42* gene is alternatively spliced to generate mRNAs with two different 3’UTRs that encode proteins with distinct C-termini. The C-termini of these Cdc42 proteins include CAAX and CCAX motifs for post-translational prenylation and palmitoylation, respectively. Palmitoyl-Cdc42 protein was previously shown to contribute to dendrite maturation, while the prenyl-Cdc42 protein contributes to axon specification and its mRNA was detected in neurites. Here, we show that the mRNA encoding prenyl-Cdc42 isoform preferentially localizes into PNS axons and this localization selectively increases *in vivo* during PNS axon regeneration. Isoform specific siRNA knockdowns, rescue experiments with siRNA-resistant Cdc42 isoforms, and pharmacologically targeting Cdc42 activity indicate that prenyl-Cdc42 promotes axon growth while the palmitoyl-Cdc42 has little growth promoting activity. The growth promotion by prenyl-Cdc42 requires axonal mRNA localization with localized translation and an intact C-terminal CaaX motif for localized prenylation of the encoded protein. Together, these data show that alternative splicing of the *CDC42* gene product generates an axon growth promoting locally synthesized prenyl-Cdc42 protein.

**SUMMARY STATEMENT:** Axon regeneration drives selective localization of alternatively spliced CDC42 isoform to PNS axons.

## INTRODUCTION

Concentrating proteins in subcellular regions is used to create unique functional domains in polarized cells. This is achieved by specifically transporting proteins from their sites of translation to subcellular regions through protein targeting sequences that are used for protein localization (Emanuelsson et al., 2007). Transport of mRNAs to subcellular domains with subsequent localized translation in those domains is also used to spatially regulate the proteome of polarized cells. For some mRNAs, this localized translation imparts unique functions to the cell. As remarkably polarized cells with widely separated and specialized subcellular domains, neurons have provided a very useful model system to profile the populations of mRNAs that are transported into axons and dendrites as well as to test functions of proteins synthesized locally in these subcellular compartments (Kar et al., 2018; Terenzio et al., 2017). Indeed, axons can extend to lengths that are more than 1000-fold longer than the neuron’s cell body diameter, and localized synthesis of new proteins in distal axons provides a level of autonomy to respond to extracellular stimuli (Sahoo et al., 2018). Both growth-promoting and growth-inhibiting stimuli regulate intra-axonal protein synthesis to support directional growth and branching of axons (Rangaraju et al., 2017; Terenzio et al., 2017).

RNA profiling studies have shown hundreds to thousands of mRNAs localized into axons of sensory, cortical, retinal ganglion cell, and motor neurons (Kar et al., 2018). These populations include several mRNAs encoding proteins that can impact the axonal cytoskeleton, either directly through generation of cytoskeletal proteins or indirectly by generation of proteins that regulate polymerization, depolymerization, severing or branching of the cytoskeleton. The mRNA encoding the small GTPase RhoA, whose activation has been shown to trigger axon retraction, localizes into and is translated within axons in response to growth inhibitory stimuli (Wu et al., 2005). RhoA belongs to the Rho GTPase family that also includes Cdc42 and Rac proteins. Rho GTPases regulate cytoskeleton organization and antagonistic functions among these family members have been reported in neurons (Etienne-Manneville and Hall, 2002). RhoA activation leads to neurite retraction through an increase in actomyosin contractility and actin depolymerization, while Cdc42 and Rac activation lead to neurite extension through promoting actin polymerization (Da Silva et al., 2003; Jalink et al., 1994; Luo, 2000; Luo et al., 1994). Intra-axonal translation of *RhoA* mRNA increases after exposure to semaphorin and chondroitin sulfate proteoglycan (CSPG) triggering growth cone repulsion and/or collapse (Walker et al., 2012a; Wu et al., 2005). The *CDC42* gene undergoes alternative splicing to generate two distinct *Cdc42* mRNAs, and recent work points to local translation of the *Cdc42* mRNA variant containing its exon 7 in neurites of cultured neurons (Ciolli Mattioli et al., 2019). We find that the exon 7 containing-Cdc42 mRNA, which encodes prenylated Cdc42 (*prenyl-Cdc42*; also known as the ‘placental isoform’ (Wirth et al., 2013)), selectively localizes into axons, and this axonally localized mRNA accounts for axon growth promoting effects of Cdc42 protein. In contrast, the *Cdc42* mRNA variant encoding the palmitoylated *Cdc42* mRNA isoform (*palm-Cdc42*; also known as the ‘brain isoform’ (Wirth et al., 2013)) has no role in axon growth and does not localize into axons. Cdc42’s contribution to axonal growth has been known for many years, and our data indicate that this is uniquely driven by the prenyl-Cdc42 isoform through axonal localization of its mRNA and its carboxyl (C)-terminal modification of its locally translated protein product.

## RESULTS

### Prenyl-Cdc42 mRNA isoform uniquely localizes into axons

The rat *CDC42* gene consists of 7 exons that generate two mRNA isoforms by alternative splicing to utilize exon six or seven (Figure 1A). The splice variant ending with exon six encodes *palm-Cdc42* mRNA that generates palmitoylated Cdc42 protein, while that mRNA skipping exon six and ending in exon seven encodes the *prenyl-Cdc42* mRNA that generates prenylated Cdc42 protein (Wirth et al., 2013). This alternative splicing to generate two *Cdc42* mRNA isoforms with unique 3’UTRs and different C-termini is conserved from fish to mammals (Supplemental Table S1). Ciolli Mattioli et al. (2019) recently reported that the *prenyl-Cdc42* mRNA variant and not the *palm-Cdc42* mRNA can localize into neurites of mouse embryonic stem cell-derived and primary cortical neurons, but they did not distinguish between dendrites and axons (Ciolli Mattioli et al., 2019). However, using RNA-Seq data for cortical neurons Taliaferro et al. (2016) show near equivalent proportions of *prenyl-Cdc42* and *palm-Cdc42* mRNAs in neurites (Supplemental Table S2). By mining published next-generation sequencing (RNA-Seq) data from axon only RNA isolates from cultured embryonic mouse sensory and motor neurons, we find that an overall higher proportion of the *prenyl-Cdc42* mRNA localizes into axons than the *palm-Cdc42* mRNA (Supplemental Table S2) (Briese et al., 2016; Minis et al., 2014). RNA-seq data from adult mouse DRG cultures similarly show higher proportion of the *prenyl-Cdc42* mRNA in axons than *palm-Cdc42* mRNA (Supplemental Table S2).

**Figure 1:**
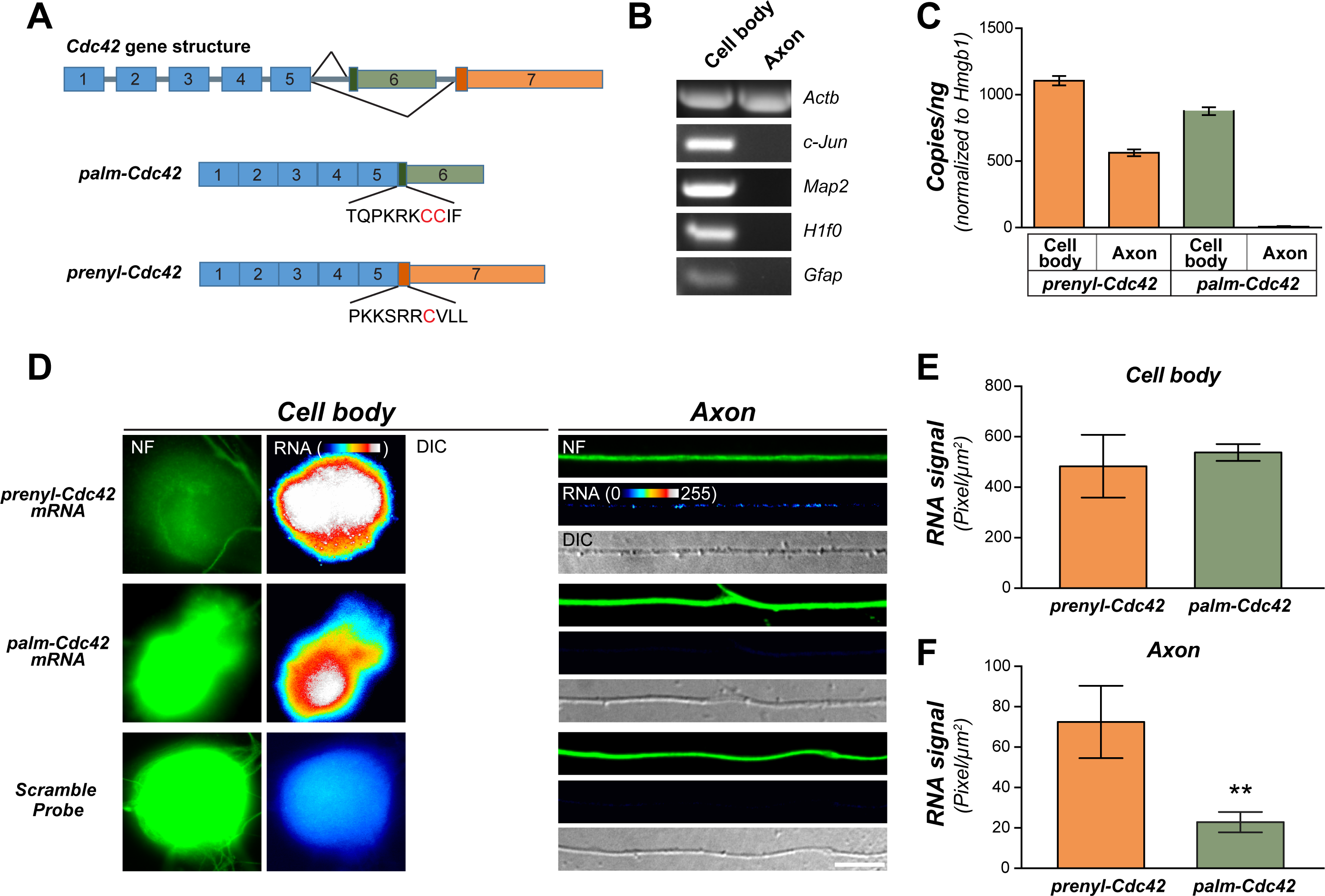
Alternatively spliced Cdc42 mRNA isoforms differentially localize to axons of sensory neurons. **A**, Schematic for alternative splicing for *Cdc42* mRNAs. The two *Cdc42* mRNAs have distinct 3’UTRs from alternative use of exons 6 or 7. These two mRNAs encode proteins with different C-termini. **B**, Representative ethidium bromide stained agarose gel of RT-PCR for RNA isolated from axonal and cell body compartments of dissociated DRG neurons. **C**, RT-ddPCR for prenyl-Cdc42 and palm-Cdc42 isoforms is shown as mean copy number ± SEM; *Hmgb1* was used for input RNA normalization (N = 3 biological replicates). **D**, Representative exposure-matched FISH/IF images for *Cdc42* mRNA isoforms FISH and neurofilament (NF) in adult DRG neuron cultures [Scale bars = 10 µm]. **E-F**, Quantification of FISH signal intensities shown as mean pixel intensity ± SEM for cell body (**E**) and axons (**F**; n ≥ 60 neurons from 3 independent cultures; ** P < 0.01 by one-way ANOVA with pair-wise comparison and Tukey post-hoc tests).

To more directly test the possibility for selective axonal localization of *prenyl-Cdc42* mRNA, we cultured L4-6 DRG neurons from adult rat sensory neurons on a porous membrane to allow for isolation of axonal RNA (Zheng et al., 2001). Reverse transcriptase-coupled polymerase chain reaction (RT-PCR) showed that the axonal preparations contained *Actb* mRNA but not the somatodendritic-restricted *Map2* mRNA, cell body-restricted *c-Jun* and *H1f0* mRNAs, or glial *Gfap* mRNA (Figure 1B). Using reverse transcription-coupled ddPCR (RT-ddPCR), comparable copy numbers for *palm*- and *prenyl-Cdc42* mRNAs were detected in the DRG cell body RNA preparations (Figure 1C). However, *prenyl-Cdc42* mRNA was much more abundant than *palm-Cdc42* mRNA in the axonal RNA isolates, with only a few copies of *palm-Cdc42* mRNA detected in the axonal RNA isolates (Figure 1C). Fluorescence *in situ* hybridization (FISH) also showed granular *prenyl-Cdc42* mRNA signals in axons of cultured DRG neurons, but axonal signals for *palm-Cdc42* mRNA were comparable to the scrambled probe control (Figure 1D-F).

With this preferential localization of *prenyl-Cdc42* mRNA into axons of cultured adult DRG neurons, we asked if *preny-Cdc42* mRNA localizes into axons *in vivo*. FISH signals for *prenyl-Cdc42* mRNA were very low in uninjured sciatic nerve axons, but signals could be seen in Schwann cells in the nerve sections (Figure 2A). However, the axonal *prenyl-Cdc42* mRNA FISH signals were increased seven days after seven days sciatic nerve crush (Figure 2A; Supplemental Figure 1A). Quantifying the FISH signals across multiple animals showed a significant increase in axonal *prenyl-Cdc42* mRNA in the regenerating axons compared to uninjured axons, while the axonal *palm-Cdc42* mRNA signals were again comparable to the scrambled control (Figures 2A-B). As a second test for *Cdc42* mRNA isoform levels in sciatic nerve axons, we used RT-ddPCR with a sciatic nerve axoplasm preparation that is highly enriched in axonal contents (Hanz et al., 2003). Again, there was a significant increase in *prenyl-Cdc42* mRNA in the regenerating nerve axoplasm of the compared to naïve nerve axoplasm with very low signals for *palm-Cdc42* mRNA in both (Figure 2C). Interestingly, DRG levels of *prenyl-Cdc42* and *palm-Cdc42* mRNA did not significantly change with injury (Figure 2C), suggesting that the increase in axonal *preny-Cdc42* mRNA in regenerating axons is driven either by increased transport into or stabilization of the mRNA in axons.

**Figure 2:**
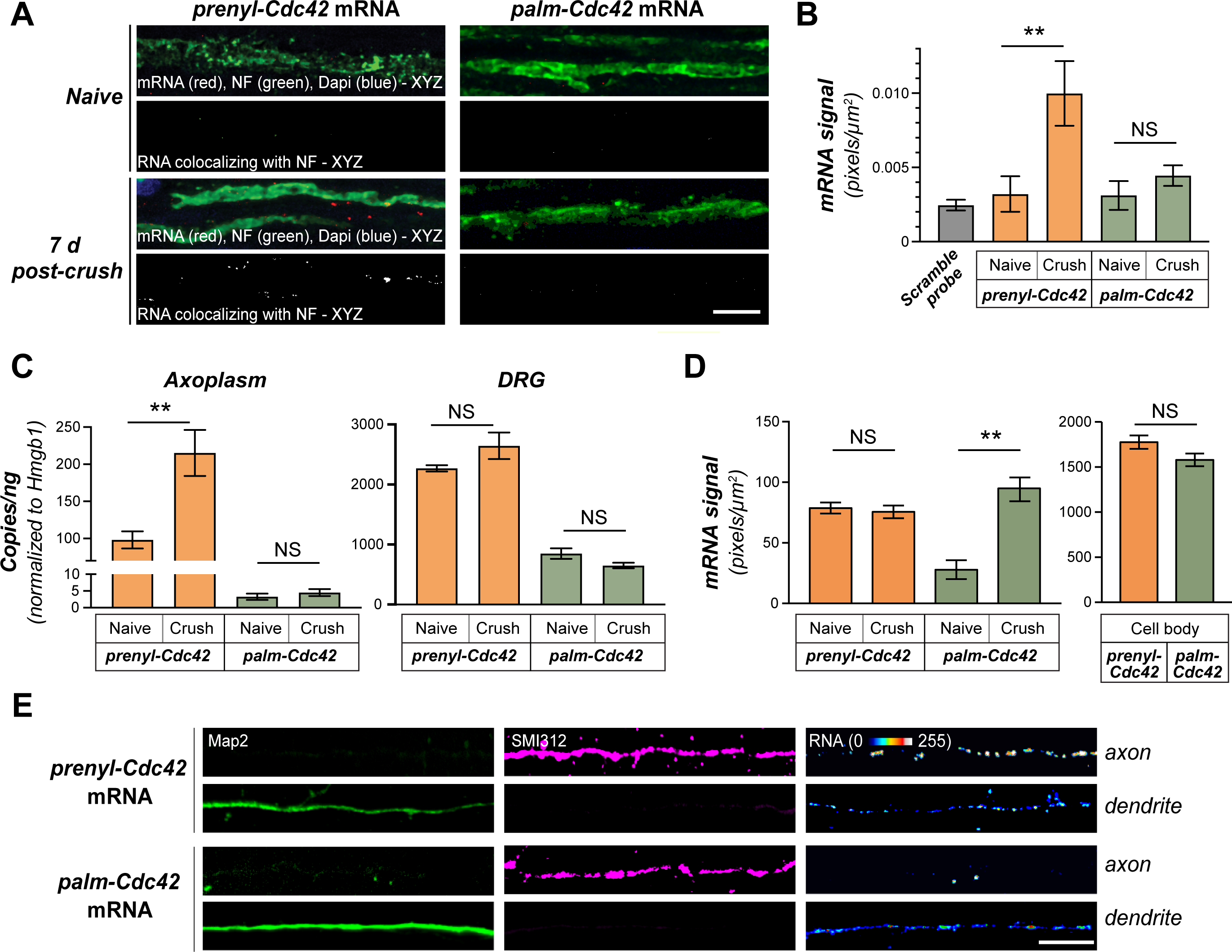
Prenyl-Cdc42 mRNA localizes to axonal processes. **A**, Representative FISH/IF images for naïve and 7 day post-crush injured sciatic nerve are shown. Upper row of each image set shows merged confocal XYZ projections; lower row of each image set shows RNA signal overlapping with NF across individual optical planes that was extracted to a separate channel and projected as XYZ images. For scrambled probe signals see Supplemental Figure S1A. **B**, Quantification of FISH signals for RNA probe signals overlapping with NF from A are shown as mean ± SEM (N = 15; ** p < 0.01, NS = non-significant, by one-way ANOVA with pair-wise comparison with Tukey post-hoc tests). **C**, RTddPCR for RNA isolated from extruded sciatic nerve axoplasm or L4-6 DRG lysates shown as mean ± SEM. mRNA copy number is normalized to *Hmgb1* mRNA levels (N = 6; ** P < 0.01 and NS = not significant by one-way ANOVA with pair-wise comparison with Tukey post-hoc tests). **D-E**, Representative exposure matched FISH/IF images of DIV 7 cortical neuron for *prenyl-Cdc42* and *palm-Cdc42* mRNAs plus phosphorylated NF (SMI312) as an axonal marker and MAP2 as a dendritic marker show in **E**. For RNA signals in corresponding cell bodies, refer to Supplemental Figure S1B. Quantification of *prenyl-Cdc42* and *palm-Cdc42* mRNA signals in axons, dendrites, and cell bodies across multiple DIV 7 cortical neuron culture preparations is shown as mean pixel intensity ± SEM in **D** (n ≥ 60 neurons in 3 independent transfections; ** P < 0.01 and NS = not significant by one-way ANOVA with pair-wise comparison with Tukey post-hoc tests) [Scale bars = 10 µm].

Since Taliaferro et al. (2016) detected *palm-Cdc42* mRNA in mouse cortical neurites by RNA-seq analyses (Supplemental Table S2), we asked if these *Cdc42* mRNA variants might show differential localization in axons and dendrites. The *palm-Cdc42* mRNA was detected in dendrites and not axons of DIV7 cortical neurons from E18 rat pups by FISH, while *prenyl-Cdc42* mRNA was detected in both axons and dendrites (Figure 2D-E; Supplemental Figure S1B). Thus, only the *prenyl-Cdc42* mRNA isoform localizes into axons, while the *palm-Cdc42* mRNA shows strictly somatodendritic localization in these DIV7 cortical neurons. This dendritic localization of *palm-Cdc42* mRNA conflicts with a recent report that *palm-Cdc42* mRNA is restricted to the cell body in mouse ES cell-derived and DIV21 mouse cortical neurons by Ciolli Mattiolli et al. (2019).

### Prenyl-Cdc42 3’UTR drives axon localization

The unique 3’UTR of *prenyl-Cdc42* was suggested to be sufficient for translation of mCherry reporter mRNA in neurites of ES cell-derived neurons by a puromycin-proximity ligation assay and immunoblotting (Ciolli Mattioli et al., 2019). We used myristoylated (MYR) eGFP reporter constructs containing the 3’UTRs of *prenyl-Cdc42* vs. *palm-Cdc42* mRNAs (GFP^MYR^3’palm-cdc42 and GFP^MYR^3’prenyl-cdc42, respectively) to compare potential axonal localizing activity of *CDC42* exons 6 vs. 7 in adult DRG cultures. By FISH for *GFP* mRNA, only the GFP^MYR^3’prenyl-cdc42 transfected neurons showed detectable axonal *GFP* RNA signals (Figure 3A-C). Axonal FISH signals for *GFP*^*MYR*^*3’palm-Cdc42* mRNA were not significantly different than those of the *GFP*^*MYR*^ mRNA carrying the 3’UTR of *Actg* mRNA that does not localize into DRG axons (Figure 3C) (Willis et al., 2007). The MYR tag of the GFP markedly limits diffusion of the protein synthesized in neurites, so that sites of protein synthesis can be visualized using FRAP (Aakalu et al., 2001; Yudin et al., 2008). FRAP studies revealed that distal axons of DRGs expressing *GFP*^*MYR*^*3’prenyl-cdc42* mRNA showed significantly higher GFP^MYR^ recovery after photobleaching than those expressing *GFP*^*MYR*^*3’palm-cdc42* (Figure 3D-E). Pre-incubation with the translation inhibitor anisomycin attenuated the post-bleach fluorescent recovery of GFP^MYR^3’prenyl-cdc42, indicating that the axonal recovery seen for this reporter derives from locally synthesized G (Figure 3D-E). The GFP^MYR^3’palm-cdc42 showed more recovery than GFP^MYR^3’γ-actin transfected neurons, which may reflect the low axonal levels of *palm-Cdc42* mRNA seen by RTddPCR in Figure 1C. Taken together, these data show that the 3’UTR from *CDC42* gene exon 7 is sufficient for axonal localization and translation of *prenyl-Cdc42* mRNA.

**Figure 3:**
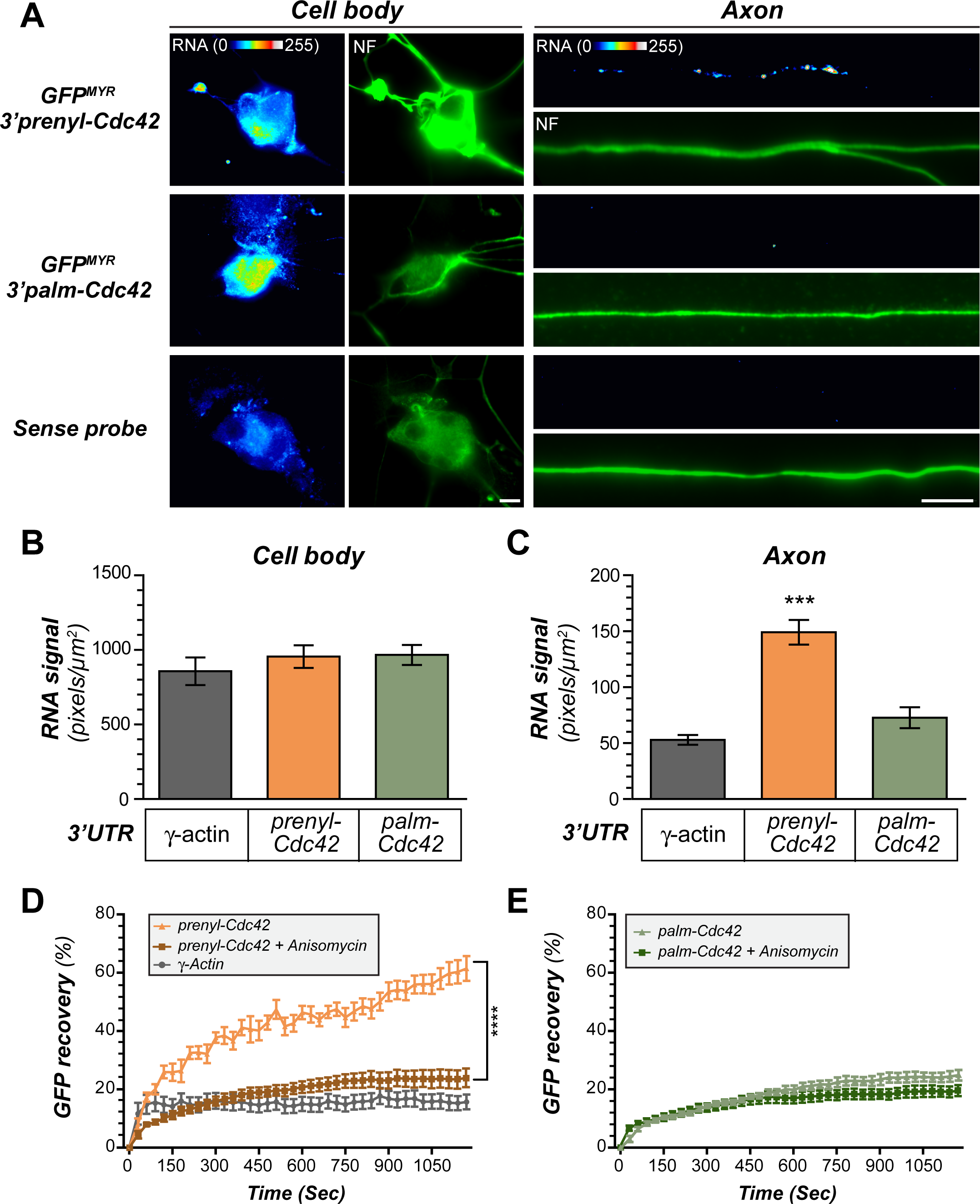
prenyl-Cdc42 mRNA isoform 3’UTR drives axonal mRNA localization and translation. **A**, Representative exposure-matched FISH/IF images for GFP mRNA and NF in DRG neurons transfected with GFP^MYR^ reporter containing 3’UTRs of *prenyl-Cdc42* or *palm-Cdc42* mRNAs (GFP^MYR^-3’prenyl-Cdc42 and GFP^MYR^3’palm-Cdc42, respectively) are shown [Scale bars = 10 µm]. **B-C**, Quantification of GFP mRNA signals from FISH for cell bodies (**B**) and axons (**C**) shown as mean pixel intensity ± SEM. GFP^MYR^3’γ-actin was used as a negative control (N ≥ 45 neurons in 3 transfections; *** p ≤ 0.001 by one-way ANOVA with pair-wise comparison and Tukey post-hoc). **D-E**, FRAP analyses for distal axons of DRG neurons expressing indicated GFP^MYR^ reporter constructs as in A are shown as mean % normalized average recovery ± SEM. The GFP^MYR^3’prenyl-Cdc42 shows significantly greater fluorescence recovery than GFP^MYR^3’palm-Cdc42 and GFP^MYR^3’γ-actin. Note that translation inhibition with anisomycin prior to photobleaching shows that the GFP^MYR^3’prenyl-Cdc42 recovery requires protein synthesis (N ≥ 15 neurons tested over 3 culture preparations; **** P < 0.001 by two-way ANOVA with pair-wise comparison and Tukey post-hoc tests).

Alternative splicing of the *CDC42* gene transcript accounts for the different 3’UTRs of the endogenous *prenyl-* and *palm-Cdc42* mRNAs. The GFP^MYR^ constructs used above would not be subjected to these splicing events, and retention of proteins at splice sites has been suggested to impact subcellular localization and translation of the endogenous mRNAs (Le Hir et al., 2016). To address the possibility that splicing rather than the RNA sequence determines axonal localization of *prenyl-Cdc42* mRNA, we generated *CDC42* ‘mini-gene’ constructs that have GFP cDNA fused to rat *CDC42* exons 4, 5 and 6 or exons 4, 5 and 7 to mimic the splicing events for *palm-Cdc42* and *prenyl-Cdc42* mRNAs, respectively (Supplemental Figure S2A). Exons 4 and 5 were separated by the endogenous 103 nucleotide (nt) intervening intron; exons 5 and 6 or 5 and 7 were separated by 500 nt intronic sequence downstream of exon 5 and plus 500 nt intronic upstream of exon 6 or 7 (i.e., 1000 nt intronic sequence). By RT-PCR DRGs transfected with these mini-gene constructs showed PCR products at the anticipated sizes of the mature (i.e., spliced) mRNAs for GFP + *CDC42* exons 4-6 and GFP + *CDC42* exons 4-5 + 7 (Supplemental Figure S2B). By FISH, GFP mRNA in cell bodies showed comparable levels for DRG neurons transfected with the mini-gene constructs (Supplemental Figure S2C). Robust axonal localization was seen for the mini-gene construct containing *CDC42* exon 7 but not for exon 6 (Supplemental Figure S2D). Neither mRNAs from the mini-gene containing *CDC42* exon 6 nor the GFP-3’palm-Cdc42 showed axonal levels significantly greater than the GFP control (Supplemental Figure S2D). These data further emphasize that 3’UTR sequences within exon 7 of the *CDC42* gene are necessary and sufficient for axonal localization, and this explains the more robust axonal localization of *prenyl-Cdc42* mRNA compared to *palm-Cdc42* mRNA.

### Axonal prenyl-Cdc42 mRNA drives axon growth

Cdc42 protein has been shown to localize to growth cones and its activity supports neurite outgrowth (Friesland et al., 2013; Matsuura et al., 2004; Myers et al., 2012). Recent work linked the prenyl-Cdc42 protein isoform to axon specification in CNS neurons while the palm-Cdc42 protein isoform was needed for dendritic spine development (Yap et al., 2016). To determine whether axonal localization of *prenyl-Cdc42* mRNA contributes to these growth effects of Cdc42 protein, we selectively knocked down prenyl-vs. palm-Cdc42 or both using isoform-specific or pan Cdc42 siRNAs (siPrenyl, siPalm, and pan-siCdc42; Figure 4A). The pan-siCdc42 depleted both *Cdc42* mRNA isoforms (Figure 4B-C) and, consistent with a previous report (Chandran et al., 2016), decreased axon growth in the DRG cultures (Figure 4D-E). siPalm depleted *palm-Cdc42* mRNA, but had no effect on *prenyl-Cdc42* mRNA levels nor did it affect axon lengths (Figure 4B-C, D-E). In contrast, the siPrenyl transfection significantly decreased axon growth to levels comparable to depletion of both mRNA isoforms using pan-siCdc42 (Figure 4D-E). siPrenyl depleted *prenyl-Cdc42* mRNA by more than 70 %, but also decreased the *palm-Cdc42* mRNA by 40-50 % (Figure 4B-C).

**Figure 4:**
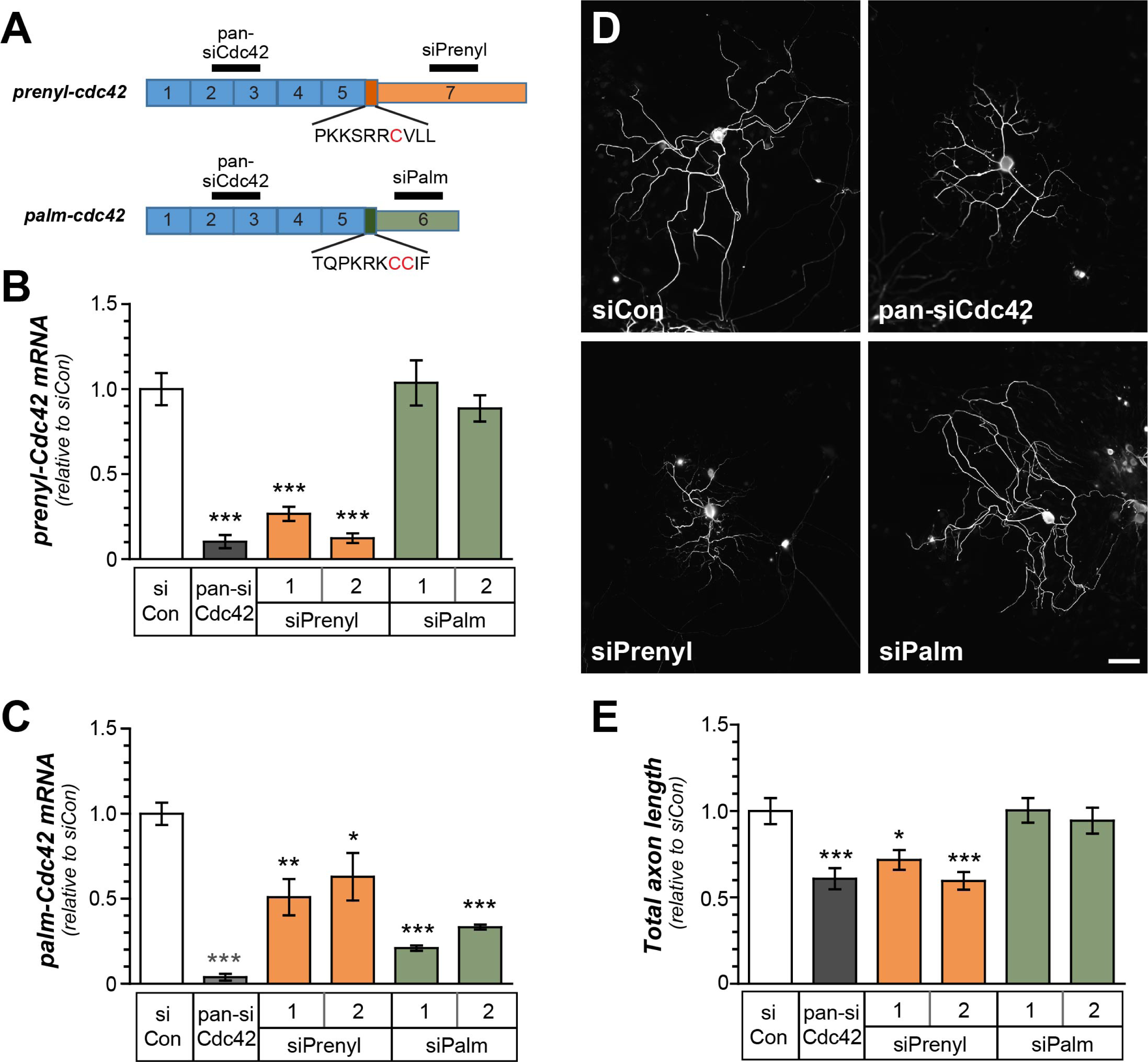
prenyl-Cdc42 isoform depletion decreases axon growth. **A**, Schematic for siRNAs targeting both *Cdc42* isoforms (pan-siCdc42) and *prenyl-Cdc42* isoform and *palm-Cdc42* mRNA isoforms. Two closely spaced siRNAs were designed to target each isoform mRNA (indicated as siPrenyl 1 and 2 and siPalm 1 and 2). **B-C**, Cdc42 mRNA levels were assessed by RT-ddPCR using primers selective for the *prenyl-Cdc42* (**B**) and *palm-Cdc42* (**C**) mRNA, where both were normalized to *Hmgb1* mRNA. Levels are shown as mean fold-change relative to control siRNA (siCon) ± SEM after (N = 3; * P < 0.05, ** P < 0.01 and *** P < 0.005 by one-way ANOVA with pair-wise comparison and Tukey post-hoc tests). **D-E**, Representative images for NF immunofluorescence in rat DRG neuron cultures transfected with siRNAs from A at 48 hours post-transfection shown in **D**. Total axon length/neuron for transfected neurons shown as mean ± SEM (N ≥ 90 neurons tested over 3 culture preparations; * P < 0.05 and *** P < 0.005 by one-way ANOVA with pair-wise comparison and Tukey post-hoc tests) [Scale bar = 100 µm].

The comparable effect of pan-siCdc42 and siPrenyl on axon outgrowth is consistent with the hypothesis that axonally localizing *prenyl-Cdc42* mRNA selectively supports axon growth, but we could not rule out contributions of the *palm-Cdc42* mRNA isoform to axon growth given that its levels were decreased by both pan-siCdc42 and siPrenyl transfections. Thus, we tested the ability of siRNA-resistant (‘si-resistant’) prenyl-Cdc42 and palm-Cdc42 cDNA constructs to rescue the decrease in axonal growth seen after pan-siCdc42 transfection (Figure 5A). Axon growth was only fully rescued by expressing the si-resistant *prenyl-Cdc42* mRNA with its axonally localizing 3’UTR (Figure 5B-C). There was no rescue when the si-resistant *prenyl-Cdc42* mRNA included the non-localizing 3’UTR of the GFP construct (Figure 5B-C). The si-resistant *palm-Cdc42* mRNA similarly did not rescue the axon growth deficit in the siCdc42-transfected DRGs (Figure 5B-C). Notably, branching of axons was not affected by knockdown of Cdc42 or rescue transfections with si-resistant prenyl-Cdc42 or palm-Cdc42 constructs (Figure 5D), indicating that Cdc42 contributes to axon elongation and not axon branching. Taken together these data indicate that the axonally localizing *prenyl-Cdc42* mRNA isoform supports axon growth in adult sensory neurons, but not the *palm-Cdc42* mRNA isoform.

**Figure 5:**
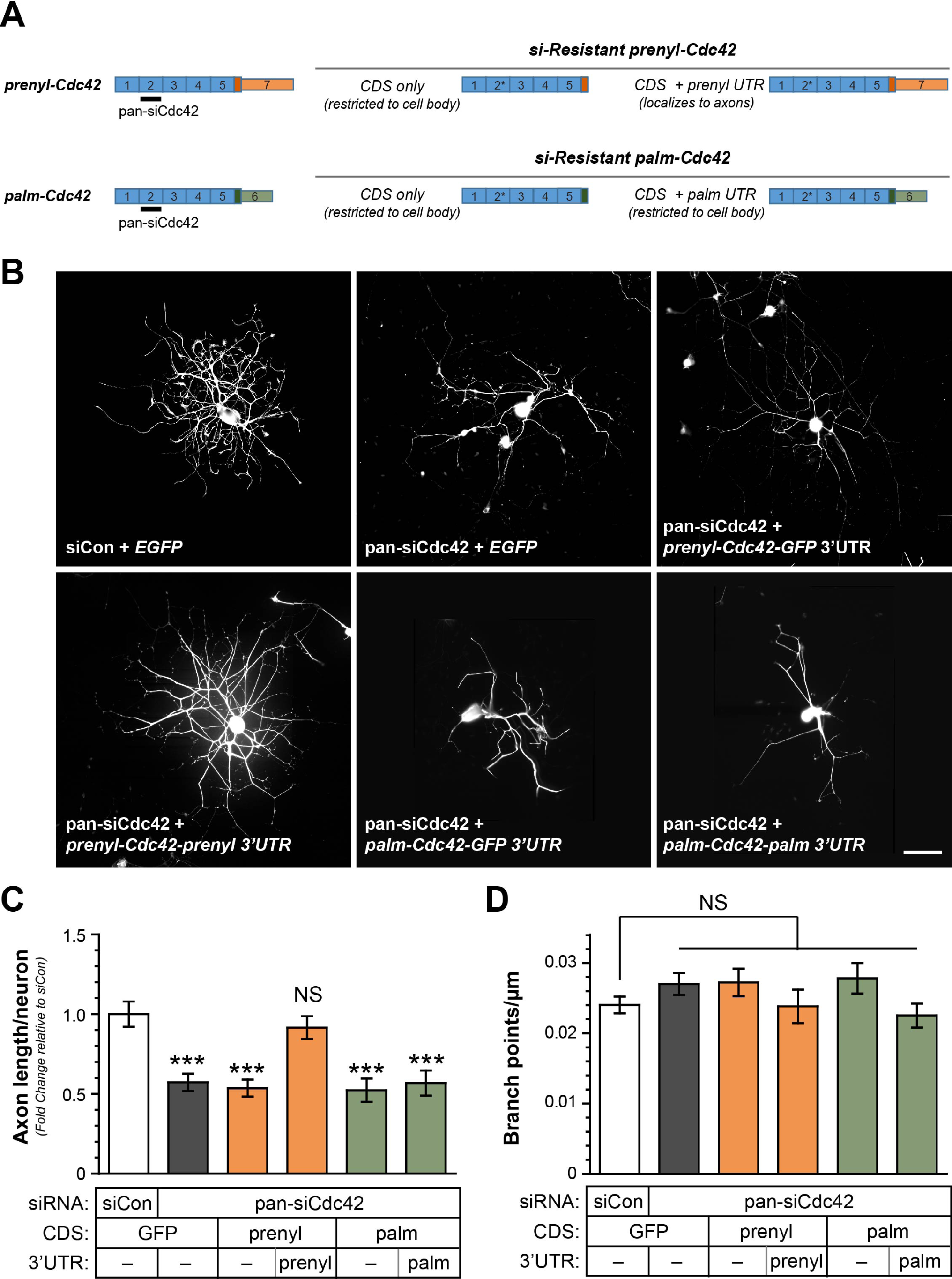
Axonal prenyl-Cdc42 mRNA rescues axon growth reduction by siRNA. **A**, Schematic for siRNA-resistant cDNA constructs used in panels B-D. **B**, Representative NF immunofluorescent images for rat DRG neurons transfected with siCon or pan-siCdc42 plus si-resistant cDNA constructs from A at 96 hours post-transfection [Scale bar = 20 µm]. **C-D**, Total axon length/neuron for DRG neurons transfected as in B is shown in **C** as mean fold-change ± SEM relative to siCon + GFP transfection. **D** shows axon branching as mean branch points/µm ± SEM (N ≥ 150 neurons over 3 replicates; *** P < 0.005 by one-way ANOVA with pair-wise comparison and Tukey post-hoc tests).

To determine if activity of prenyl-Cdc42 protein is needed for axon growth, we tested two small molecules that directly and specifically target Cdc42 activity. ZCL278 and ZCL367 were identified by an *in silico* screen for chemical structures predicted to bind to residues in Cdc42 protein that interact with the Cdc42-specific guanine nucleotide exchange factor (GEF) Intersectin (Friesland et al., 2013). ZCL278 has dual activities in that it decreases Cdc42 activity *in vitro* and in cultured cells when Cdc42 is ligand-activated (Friesland et al., 2013), while it can increase Cdc42 activity when the protein is not ligand-activated (Aguilar et al., 2019). Consistent with this, axon growth was significantly increased by 10 and 25 µM ZCL278 treatment, while axon growth was significantly attenuated by 25 µm ZCL367 treatment (Supplemental Figure S3A). The growth-promoting effect of ZCL278 in these adult DRG cultures was blocked by transfection with siPrenyl or pan-siCdc42, but DRG neurons transfected with siPalm showed growth promotion by ZCL278 similar to siCon transfected DRG neurons (Supplemental Figure S3B). In contrast, axon growth was equally decreased in siCon, pan-siCdc42, siPrenyl, and siPalm transfected neurons (Supplemental Figure S3B). Taken together with the siRNA rescue data above, these observations indicate that activity of the prenyl-Cdc42 and not the palm-Cdc42 isoform promotes axon growth.

### Axonal RNA localization and prenylation motif of prenyl-Cdc42 mRNA are needed for optimal axonal outgrowth

Although only differing by 11 amino acids, unique functions for neuronal prenyl-vs. palm-Cdc42 protein isoforms have been found in previous studies (Kang et al., 2008; Yap et al., 2016). We wondered if distinct subcellular localization of *prenyl-* and *palm-Cdc42* mRNAs in the DRG neurons contributes to the unique functions of prenyl- and palm-Cdc42 protein isoforms. So we asked if targeting the *palm-Cdc42* mRNA into axons would affect axon growth. For this, we generated si-resistant palm-Cdc42 expressing cDNAs where the *palm-Cdc42* mRNA 3’UTR was replaced by the axonal-targeting *prenyl-Cdc42* mRNA 3’UTR (Figure 6A). pan-siCdc42 transfected DRG neurons showed partial rescue of axon growth with expression of the axonally targeted, si-resistant *palm-Cdc42* mRNA (Figure 6B). This observation suggests that either Cdc42 isoform can support axon growth, but notably a full rescue effect required axonally targeted *prenyl-Cdc42*.

**Figure 6:**
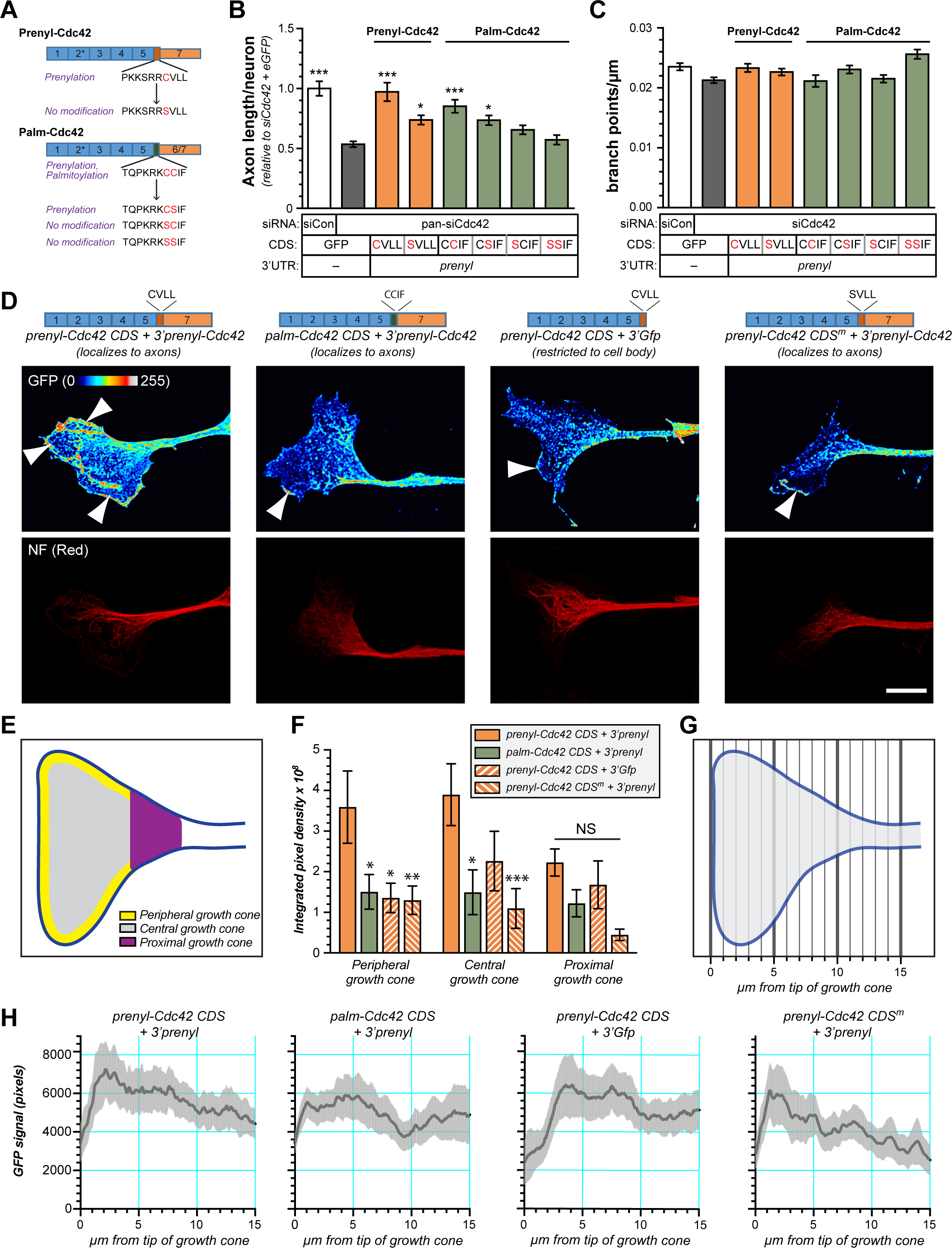
C-terminal CaaX motif is needed for optimal axon growth by axonal prenyl-Cdc42 mRNA. **A**, Schematic for si-resistant Cdc42 isoforms with CaaX and CCaX motifs. **B-C**, Total axon length/neuron for DRG neurons transfected with pan-siCdc42 plus the indicated si-resistant CaaX and CCaX mutants for Cdc42 isoforms is shown in **B** as mean fold change relative to siCon + eGFP ± SEM. **C** shows axon branching as mean branch points/µm axon length ± SEM. Note that mRNAs for all Cdc42 mutants were targeted into axons using the 3’UTR from *prenyl-Cdc42*. (N ≥ 120 neurons over 3 transfections; * P < 0.05 and *** P < 0.005 by one-way ANOVA with pair-wise comparison and Tukey post-hoc). **D**, Representative exposure-matched fluorescent images for GFP (upper row) and NF (lower row) for indicated GFP-Cdc42 expression constructs with mutated C-termini in growth cones of DRG neuron are shown as maximum XYZ projection from deconvolved confocal Z stacks. mRNAs for the cDNA constructs were cell body-restricted using 3’UTR of *GFP* or axonally localizing using 3’UTR of *prenyl-Cdc42* mRNAs. Representative IF images in each column are for neurons expressing the indicated Cdc42 cDNA construct. [Scale bar = 10 µm]. **E-F**, Schematic peripheral, central and proximal domains of the growth cone used for GFP-Cdc42 signal quantitation across the growth cone shown in **E**. Quantitation of GFP-Cdc42 intensity in growth cone domains defined in E is shown as mean ± SEM in **F**. The protein product of the axonally localizing *GFP-Prenyl-Cdc42 CDS + 3’prenyl-cdc42* shows significantly greater signals in the periphery and central domains compared to protein products of cell body restricted *GFP-Prenyl-Cdc42 CDS + 3’gfp* and axonally localizing *GFP-Palm-Cdc42 CDS + 3’prenyl-cdc42* and *GFP-Prenyl-Cdc42 CDS SVLL mutant + 3’prenyl-cdc42* (N ≥ 9 growth cones over 3 biological replicates; * P < 0.05, ** P < 0.01, and *** P < 0.001 by two-way ANOVA with Tukey post-hoc) **G-H**, Schematic of method for unbiased assessment of GFP-Cdc42 signals across growth cone shown in **G**. Quantitation of growth cone signals for indicated GFP-Cdc42 variants using the method defined in G is shown in **H**. The distribution of protein encoded by *GFP-Cdc42-3’prenyl* is significantly different than those from the other prenyl-Cdc42 and palm-Cdc42 constructs to *SVLL* (N ≥ 11 growth cones over 3 biological replicates; comparing *GFP-Cdc42-3’prenyl* curve to others by Kolmogorov-Smirnov test shows P < 0.0001 for all and *GFP-Cdc42-3’prenyl* has D value = 0.450 vs. *GFP-Palm-Cdc42 CDS + 3’prenyl-cdc42*, D = 0.322 vs. *GFP-Prenyl-Cdc42 CDS + 3’gfp*, D = 0.705 vs. *GFP-Prenyl-Cdc42 CDS SVLL mutant + 3’prenyl-cdc42* where lower D values indicate greater differences between curves).

With different degrees of rescue by axonally targeted *prenyl-*vs. *palm-Cdc42* mRNAs, we next asked whether the post-translational prenylation or palmitoylation contributes to the growth promoting effects of the axonally targeted *prenyl-Cdc42* mRNA. The first 180 amino acids of palm-Cdc42 and prenyl-Cdc42 proteins are identical, with the distinct C-terminal 11 amino acids coming from exons 6 and 7, respectively (Chen et al., 2012). The four C-terminal residues of these two Cdc42 isoforms contain CCaX and CaaX motifs for palmitoylation and prenylation, respectively (where C represents cysteine, a is an aliphatic amino acid and X is any amino acid) (Wirth et al., 2013). The cysteines in the CaaX and CCaX motifs are covalently modified by prenylation and palmitoylation (Nishimura and Linder, 2013), and mutating the cysteine residues to serine has been used to prevent prenylation and palmitoylation of proteins (Wirth et al., 2013). The CCaX motif in the *palm-Cdc42* protein product (CCIF) can undergo an initial prenylation at the first cysteine followed by palmitoylation at the second cysteine (Wirth et al., 2013). Thus, we generated siRNA-resistant mutants expressing palm-Cdc42 with its C-terminal CCIF residues mutated to CSIF, SCIF, and SSIF. For prenyl-Cdc42, we generated siRNA-resistant mutants with C-terminal CVLL residues mutated to SVLL (Figure 6A). Both the *prenyl-Cdc42* and *palm-Cdc42* mRNAs were targeted into axons using the 3’UTR of *prenyl-Cdc42* mRNA. The axonally targeted, siRNA-resistant *palm-Cdc42* mRNA with SCIF and SSIF mutations did not significantly increase axon growth in the pan-siCdc42 transfected neurons (Figure 6B). Axonally-targeted siRNA-resistant *palm-Cdc42* mRNA with intact CCIF motif, and to a lesser extent with CSIF, showed partial rescue of axon growth in pan-siCdc42 transfected DRG cultures compared to axonally targeted, siRNA-resistant *prenyl-Cdc42* with intact CVLL motif (Figure 6B). Notably, axon branching did not show any significant change by expression of the either wild type or these mutant Cdc42 constructs (Figure 6C). Taken together, these data suggest that both axonal mRNA localization and the intact C-terminal CVLL sequence (i.e., CaaX motif) for prenylation are needed for the full axon growth-promoting effects of the prenyl-Cdc42 isoform, with the palm-Cdc42 isoform only partially supporting axon growth when its mRNA is targeted into axons.

### Prenyl-Cdc42 and Palm-Cdc42 proteins show distinct localizations in growth cones

Since axonally targeted *palm-Cdc42* mRNA with intact CCIF motif did not promote axon growth to the same extent as *prenyl-Cdc42* mRNA, we asked if prenyl-Cdc42 and palm-Cdc42 proteins might show different localization in growth cones. We used confocal microscopy to visualize GFP-tagged prenyl-Cdc42 and palm-Cdc42 proteins in distal axons of DRG neurons transfected with Cdc42 constructs with or without axonally localizing 3’UTR from *CDC42* exon 7 (Figure 6D). The protein product of the axonally targeted *GFP-prenyl-Cdc42* mRNA showed fluorescent signals concentrated at the periphery of the growth cone, frequently with localization at or just beneath the cytoplasmic membrane at the growth cone’s leading edge (Figure 6D). In contrast, signals from axonally targeted translation products for *GFP-prenyl-Cdc42* mRNA with CVLL to SVLL mutation to prevent prenylation and axonally targeted translation products for *GFP-palm-Cdc42* mRNA rarely concentrated signals at the leading edge of the growth cone (Figure 6D). Replacing the axonal-targeting 3’UTR of *GFP-prenyl-Cdc42* with a cell body-restricting 3’UTR gave signals that rarely concentrated at the growth cone’s leading edge (Figure 6D). Quantifying the signal intensities for these GFP-prenyl-Cdc42 and GFP-palm-Cdc42 showed significantly higher levels of protein at the growth cone periphery for the axonally targeted *GFP-prenyl-Cdc42*-*3’prenyl-Cdc42* than cell body-restricted *GFP-prenyl-Cdc42-3’gfp* or axonally targeted *GFP-palm-Cdc42-3’prenyl-Cdc42* (Figure 6E-H). The protein encoded by *GFP-prenyl-Cdc42-3’prenyl-Cdc42* with SVLL mutation showed growth cone peripheral signals intermediate between axonally targeted *GFP-prenyl-Cdc42*-*3’prenyl-Cdc42* and the cell body-restricted *GFP-palm-Cdc42-3’gfp*. This suggests that axonal localization of *Prenyl-Cdc42* mRNA is needed for targeting the protein to the growth cone periphery.

## DISCUSSION

Intra-axonally synthesized proteins contribute to axon growth, both during development and regeneration, and a few thousand mRNAs are now known to localize into axons *in vitro* (Kar et al., 2018). Our data indicate that the *prenyl-Cdc42* mRNA, and not the *palm-Cdc42* isoform, is transported into axons where its protein product promotes axon growth. With roles in actin polymerization and filopodia formation, Cdc42 activity has long been linked to axon growth and cell migration (Aguilar et al., 2017; Hall and Lalli, 2010). The *palm-Cdc42* and *prenyl-Cdc42* mRNA isoforms are generated by alternative splicing with differential inclusion of exons 6 and 7 in their mRNAs, respectively (Chen et al., 2012). Work in cultured CNS neurons has pointed to roles for the palm-Cdc42 in dendrite maturation and prenyl-Cdc42 in axon specification, respectively (Yap et al., 2016). Our data emphasize that axonal mRNA targeting by *CDC42* gene exon 7 and intact CaaX motif for prenylation are responsible for the Cdc42 protein’s axon growth promoting functions and localization to the leading edge of growth cones.

Alternative RNA splicing can increase the number of different proteins generated from a single gene. Subcellular localization of mRNAs is most frequently driven by 3’UTR sequences (Andreassi and Riccio, 2009; Gomes et al., 2014), as we show here for *prenyl-Cdc42* mRNA. *Importin β1, RanBP1, Stat3α* and *BDNF* mRNA isoforms with long 3’UTRs are selectively transported into neuronal processes (An et al., 2008; Ben-Yaakov et al., 2012; Perry et al., 2012; Yudin et al., 2008). These mRNA variants are generated by alternative poly-adenylation site usage, with an RNA localization motif introduced from the additional 3’UTR sequences. The Kuruvilla lab recently showed that an mRNA isoform for another RhoA family member, Rac1, localizes into sympathetic axons via a long 3’UTR similarly generated by alternative poly-adenylation site usage (Scott-Solomon and Kuruvilla, 2020). Similar to the alternative splicing used for *CDC42*, RNA-seq studies from neurites of neural cell lines recently showed prevalence for distal alternative last exons and additional exon usage over differential poly-adenylation site usage for subcellularly targeted mRNA isoforms (Taliaferro et al., 2016). This prevalence was similarly seen for localizing mRNA isoforms in neurites from cultures of primary cortical neurons as well as axons from embryonic DRG neuron cultures. Moreover, Taliaferro et al. (2016) observed a shift of alternative splicing to favor mRNA isoforms with localizing 3’UTRs during neuronal differentiation of CAD cells (Taliaferro et al., 2016). Although inclusion of *CDC42* exon 7 rather than exon 6 selectively generates a *Cdc42* mRNA isoform with axonally localizing 3’UTR, Cdc42 expression shifts from generating predominantly *prenyl-Cdc42* mRNA to predominantly *palm-Cdc42* mRNA during brain development (Makeyev et al., 2007). Interestingly, cultured embryonic motor and sensory as well as adult sensory neurons show overall higher levels of *prenyl-Cdc42* than *palm-Cdc42* mRNAs (Briese et al., 2016; Minis et al., 2014), and our data indicate that *prenyl-Cdc42* mRNA, and not *palm-Cdc42* mRNA, localizes into axons of cortical neurons. In contrast to mature brain, these cultured neurons continuously extend axons, suggesting that continued use of *CDC42* exon 7 drives axon growth by generating *prenyl-Cdc42* mRNA for localization into growing axons. Consistent with this notion, the axonal localization of *prenyl-Cdc42* mRNA seen here is linked to axon growth since *prenyl*-*Cdc42* mRNA levels were much lower in uninjured, non-growing axons *in vivo*.

Neuronal Cdc42 has been shown to contribute to axon growth initiation, elongation and growth cone guidance (Hall and Lalli, 2010). Linking Cdc42 functions to axon growth was determined in part based on overexpression of constitutively active or dominant negative mutations of Cdc42 (Banzai et al., 2000; Nishimura et al., 2005). Exogenous overexpression of one Rho GTPase was later shown to compete with endogenous Rho GTPases for binding to RhoGDI1, which decreased levels of endogenous Rho GTPases (Boulter et al., 2010). This complicates interpretation of previous findings for expression of the dominant negative and constitutively active Cdc42s, and emphasizes the utility of the knockdown, pharmacological Cdc42 manipulations, and rescue approaches used here for distinguishing functions of the Cdc42 isoforms. Our data further emphasize that ZCL278 can function as a Cdc42 agonist under some conditions as shown by (Aguilar et al., 2019). Since the growth promotion by ZCL278 was completely abrogated by depletion of *prenyl-Cdc42* and not *palm-Cdc42* mRNA, activity of prenyl-Cdc42 and not palm-Cdc42 is needed for axon growth promotion. Using GFP-fusion proteins, we see that Cdc42 proteins derived from the cell body-restricted *palm-Cdc42* mRNA and *prenyl-Cdc42* mRNA are transported into distal axons. It is quite likely that roles previously ascribed to Cdc42 in axon specification and growth cone guidance are from the axonally translated *prenyl-Cdc42* mRNA. Notably, previous work with overexpression of Cdc42 and mutant Cdc42 proteins did not consider axonal localization of the mRNA (Banzai et al., 2000; Nishimura et al., 2005), so results from these previous works must be interpreted in the context of differentiating cell body- and axonally-derived prenyl-Cdc42 protein.

With isoform specific overexpression and knockdown/knockout approaches, a previous report showed that prenyl-Cdc42 is required for axon development and palm-Cdc42 is required for maturation of dendritic spines (Yap et al., 2016). This implies that neuronal polarity is determined in part by the differential localization of the two Cdc42 isoforms. Though recent work in mouse ESC-derived and DIV21 primary cortical neurons only detected *Prenyl-Cdc42* mRNA in neurites (Ciolli Mattioli et al., 2019), RNA-seq data from primary cortical neuron cultures detected *palm-Cdc42* mRNA in neurites (Taliaferro et al., 2016) and we clearly show that *palm-Cdc42* mRNA selectively localizes into dendrites of DIV7 rat cortical neurons. siRNA-based depletion of both Cdc42 isoforms from hippocampal neuron cultures at DIV12 decreased dendritic spine numbers and this was rescued by expression of either prenyl- or palm-Cdc42 (Wirth et al., 2013). It is not clear if 3’UTRs for either the *prenyl-Cdc42* or *palm-Cdc42* mRNAs were included in the expression constructs used by Yap et al. (2016) and Wirth et al. (2013). Thus, determining how or if locally synthesized *prenyl-Cdc42* and/or *palm-Cdc42* mRNAs contribute to dendritic growth and maturation will be of high interest. The discrepancies for *palm-Cdc42* mRNA localization outlined above may reflect that Ciolli Mattioli et al. (2019) used DIV21 cortical cultures, which is well beyond the period of active polarization that our DIV7 were undergoing at the time of analyses.

Prenylation and palmitoylation modifications of Cdc42 proteins increase protein hydrophobicity and facilitate association with cellular membranes (Roberts et al., 2008). Palm-Cdc42 and prenyl-Cdc42 proteins have been shown to have different motility in cell membranes (Wirth et al., 2013), so they could be targeted to different membrane domains. Consistent with this, the axonally translated prenyl-Cdc42 protein showed increased localization to the periphery of growth cones compared to the more central locations for cell body synthesized prenyl-Cdc42 and palm-Cdc42 encoded by engineered mRNA with axonally targeting 3’UTR of *prenyl-Cdc42*. An intact CaaX motif for prenylation of Cdc42 was needed for the full growth promoting effects of the axonally targeted *prenyl-Cdc42* mRNA and the GFP-Prenyl-Cdc42 CVLL to SVLL mutant showed growth cone localization intermediate between the GFP fusion proteins derived from the axonally-targeted and cell body-restricted *GFP-Prenyl-Cdc42* mRNAs. This suggests that axonally translated proteins can be locally prenylated, but also that this post-translational modification either increases membrane localization of or stabilizes the protein or both. The prenyl transferases, farnesyl transferase (FT) and geranylgeranyl transferase (GGT), catalyze addition of either farnesyl or geranylgeranyl isoprenoid lipids to cysteine in the CaaX motif. Both prenyl-Cdc42 and RhoA, which is also translated locally in axons (Wu et al., 2005), are substrates for GGT and not FT (Wirth et al., 2013). Based on previous publications, both CCIF and CSIF motifs included in the axonally targeted *Palm-Cdc42* mRNA would be substrates for GGT, and hence prenylation (Nishimura and Linder, 2013; Wirth et al., 2013). Thus, our mutation data suggest that palm-Cdc42 targeted into axons through *prenyl-Cdc42* 3’UTR could undergo prenylation as the axonal growth deficit was partially rescued by axonally targeted *palm-Cdc42* mRNA with intact CCIF but not with the SCIF or SSIF mutations. Furthermore, the growth promoting effects of *prenyl-Cdc42* mRNA were lost when the mRNA restricted to the soma by replacing the 3’UTR with that of GFP.

In summary, our data indicate that the prenyl-Cdc42 isoform specifically drives Cdc42’s axon growth promoting effects in adult sensory neurons though axonal translation of its mRNA. This growth promotion also requires an intact CaaX motif for post-translational prenylation of the axonally synthesized Cdc42 protein. On first glance, our growth data seem to contradict a study that used prenylation inhibitors to link the post-translational modification to slowed axon growth on myelin-associated glycoprotein (MAG), a substrate that inhibits axon growth (Li et al., 2016). In cultured neurons, highly selective FT inhibitors were more effective than GGT inhibitors for increasing axon growth (Li et al., 2016), but the combination of the two synergistically block prenylation (Roberts et al., 2008). RhoA shows altered subcellular localization upon inhibition of GGT (Roberts et al., 2008), and in contrast to the growth promoting effects of prenyl-Cdc42 shown here, RhoA attenuates axon growth on non-permissive substrates like MAG and the protein is synthesized in axons (Walker et al., 2012b; Wu et al., 2005). Previous studies have shown compensatory geranylgeranylation through GGT for some farnesylated GTPases when FT is inhibited (Roberts et al., 2008), which could account for the increased efficacy of FT plus GGT inhibition seen by Li et al. (2016). Scott-Solomon and Kuruvilla (2020) very recently showed that sympathetic axons have GGT activity, with nerve growth factor increasing axonal GGT activity; furthermore, knocking out GGT from sympathetic neurons attenuated TrkA signaling through loss of prenylated-Rac1 from those axons. In contrast to the sympathetic neurons, the adult DRG neurons used here do not require exogenous trophic support for survival or axon growth, and our data indicate that axonal targeting of *prenyl-Cdc42* mRNA is increased during axon growth and the locally synthesized protein product increases axon growth.

## MATERIALS AND METHODS

### Animal use and neuron cultures

All animal experiments were conducted under IACUC approved protocols. For dorsal root ganglia (DRG) cultures, L4-6 ganglia were isolated from adult male Sprague Dawley rats (175-250 g) and dissociated with collagenase (Invitrogen). Cells were washed and then plated with DMEM/F12 (Cellgro) supplemented with 10% fetal bovine serum (Gemini Bio), 10 µM cytosine arabinoside (Sigma) and N1 supplement (Sigma). Cells were cultured on either glass coverslips or polyethylene-tetrathalate (PET) membrane (1 µm pores; Corning) inserts coated with poly-L-lysine (Sigma) and laminin (Millipore).

DRG cultures were transfected with 2-3 μg of plasmid using *Amaxa Nucleofection* device (Lonza) with *Basic Neuron SCN Nucleofector* kit (Program SCN-8) before plating and maintained for 48-72 hours. For depletion experiments, synthetic siRNAs (predesigned from Integrated DNA Tech., 100 nM) were transfected into DRG neurons using *DharmaFECT 3* and incubated for 3 DIV. For control, identical amounts of non-targeting siControl pools were used. RT-ddPCR was used to test the efficiency of depletion (see below). siRNA sequences were: siPrenyl (#1), 5’ GUGUUGUCAUCAUACUAAAAGCAAU 3’; siPrenyl (#2), 5’ GCAAUGUUUAAAUCAAACUAAAGAU 3’; siPalm (#1), 5’ GUGAUCAGUAGUCACAUUAGACUUG 3’; siPalm (#2), 5’ UCAGUAGUCACAUUAGACUUGUUUA 3’; siCdc42, 5’ UGGUAAAACAUGUCUCCUG 3’.

DRG cultures were exposed to the Cdc42 targeting small molecules, ZCL278 and ZCL367, after the dissociated cells were adhered to coverslips (3-4 hours post plating), and then assayed for axon growth 48 hours later. Chemical structures and rationale for selection of ZCL278 and ZCL367 as Cdc42 targeting agents was previously described (Friesland et al., 2013).

For cultures of cortical neurons, cortices from E18 male and female rat pups were harvested in Hibernate E (BrainBits) and dissociated using the Neural Tissue Dissociation kit (Miltenyi Biotec). Briefly, the harvested cortices were incubated in pre-warmed enzyme mix at 37°C for 15 min and triturated using fire-polished glass pipettes and applied to a 40 µm cell strainer. After washing and centrifugation, neurons were seeded onto poly-D-lysine coated glass coverslips at a density of 10,000 cells/coverslip in culture media containing NbActive-1 medium (BrainBits) supplemented with 100 U/ml of Penicillin-Streptomycin (Life Technologies), 2 mM L-Glutamine (Life Technologies), and 1 X N21 supplement (R&D Systems) and grown at 37°C, 5% CO_2_ for 7 days.

### Plasmid constructs

All fluorescent reporter constructs for analyses of RNA translation were based on eGFP with myristoylation element (GFP^MYR^; originally provided by Dr. Erin Schuman, Max-Plank Inst., Frankfurt) (Aakalu et al., 2001). To isolate the cDNA of rat *Cdc42* (NCBI ID: XM_008764286 for *prenyl-Cdc42*; XM_008764287 for *palm-Cdc42*), total RNA from rat DRG was reverse transcribed with *Superscript II* (Invitrogen) and then amplified by PCR using *PrimeStar HS* polymerase (Takara). Primers for amplifying the *prenyl-Cdc42* coding sequence and 3’UTR were engineered to contain terminal Not I and Sal I restriction sites. Using these restriction sites, the amplicons were then inserted into pEGFP-C3 (Clontech) vectors. *Quikchange site directed mutagenesis kit* (Stratagene) was used to introduce point mutations of CaaX or CCaX motifs.

### Fluorescence in situ hybridization (FISH) and immunofluorescence

Myristoylated EGFP was fused with 3’UTRs of Cdc42 isoforms to test their axonal localization. Transfected DRG neurons were fixed 15 min in 4 % paraformaldehyde (PFA) and hybridized with DIG-labeled GFP cRNA probes (Roche) as described previously (Merianda et al., 2013). Sheep anti-DIG (1:100, Roche) was used to detect probes with Cy5-conjugated donkey anti-sheep (1:200, Jackson ImmunoResearch). Endogenous Cdc42 isoforms were detected with 5’ Cy5-labelled Stellaris probes to the exons 6 or 7 (probe sequences available upon request; BioSearch Tech.). Hybridization conditions for Stellaris probes for cultured neurons and for tissue sections was performed as recently described (Terenzio et al., 2017).

For immunofluorescence, neurons plated on glass coverslips were fixed with 4% paraformaldehyde in PBS for 15 min at room temperature. Cultures were permeabilized with 0.3% Triton X-100 in PBS for 15 min and incubated with primary antibodies for overnight in humidified chambers at 4°C. Chicken anti-NF (1:1,000, Aves labs), SMI312 mouse anti-phospho-NF (1:500, Biolegend), and chick anti-MAP2 (1:500; Abcam) were as primary antibodies; FITC-conjugated donkey anti-chicken and donkey anti-mouse (1:200, Jackson ImmunoResearch) were used as secondary antibodies. For visualization of GFP-tagged Cdc42 proteins in growth cones, rabbit anti-GFP (1:500, Abcam) and chicken anti-NF (1:1,000, Aves labs) was used for primary antibodies and FITC-conjugated donkey anti-Rabbit (1:200, Jackson ImmunoResearch) and Cy5-conjugated donkey anti-chicken (1:200, Jackson ImmunoResearch) were used as secondary antibodies. Coverslips were washed with PBS and then incubated with secondary antibodies for an hour before mounting with Prolong anti-fade mounting solution (Invitrogen).

For analyses of FISH signals, images were obtained using an epifluorescence microscope equipped with ORCA ER CCD camera (Hamamatsu) using matched acquisition parameters. Pixel intensities were measured from distal axons from more than 50 neurons in three replicate experiments. GFP signals in the growth cone were detected by Leica SP8X confocal microscope with HyD detectors. Z-stack images were post-processed by Huygens deconvolution (Scientific Volume Imaging) integrated into the Leica LASX software (*HyVolution*). Deconvolved image stacks were projected into single plane images using the maximum pixel intensities.

‘Stellaris’ probes were used for mRNA detection in sciatic nerve sections as described (Spillane et al., 2013). Briefly, sciatic nerve segments were fixed overnight in 2% PFA at 4°C and then cryoprotected overnight in 30% sucrose at 4°C. 25 µm thick cryosections were prepared and stored at - 20°C until use. Slides were dried at 37°C for 1 hour then brought to room temperature, and all subsequent steps were performed at room temperature unless indicated otherwise. Warmed tissue sections were washed 10 min in PBS once, then 10 min in 20 mM Glycine three times. Sections were then incubated three times for 5 min in fresh 0.25M NaBH_4_. Sections were quickly rinsed in 0.1M Triethanolamine (TEA), incubated for 10 min in 0.25% Acetic Anhydride in 0.1 M TEA, washed twice with 2X saline-sodium citrate (SSC) buffer, dehydrated in graded ethanol solutions (70%, 95%, and 100% for 3 min each), and delipidated in chloroform for 5 min. After incubation in 100% and 95% ethanol for 3 min each, sections were equilibrated in 2X SSC and then incubated at 37°C in a humidified chamber in hybridization buffer without probes for 5 min before incubating overnight with probe (7 µM) and RT97 anti-NF (1:100) in hybridization buffer (Perry et al., 2016). Sections were then washed twice in 2X SSC + 10% formamide at 37°C for 30 min and once in 2X SSC for 5 min, permeabilized in PBS + 1% Triton-X100 (PBST) for 5 min, and then incubated for one hour in donkey anti-mouse FITC antibody (1:200) in 0.3% Triton-X100 supplemented with 1X blocking buffer (Roche). After washing with PBS for 5 min, sections were post-fixed in buffered 2% PFA for 15 min (Spillane et al., 2013), washed in PBS three times for 5 min, rinsed in DEPC-treated water, and mounted using *Prolong Gold Antifade*.

### RNA isolation and PCR analyses

RNA was isolated from dissociated DRG neurons or cell body/axon compartments collected from insert cultures using *RNeasy micro isolation* kit (Qiagen). Purified RNAs were quantified with *Ribogreen* (Invitrogen) and 10-50 ng of RNA were used for reverse transcription with *SensiFAST cDNA synthesis kit* (Bioline) according to the manufacturer’s protocol. To assess the purity of axonal RNA, RT-PCR was performed with primers designed to detect cell body-restricted mRNAs (*c-Jun, H1f0, Map2*) and glial cell-specific mRNAs (*Gfap*). Droplet digital PCR (ddPCR) was performed according to manufacturer’s procedure (BioRad) with either *Evagreen* (Biorad) or *Taqman* probe assays (Integrated DNA Tech). *Hmgb1* mRNA levels were used for normalizing RNA yields across different isolates; we have previously shown that cell body and axonal RNA levels of *Hmgb1* mRNA do not change after axotomy (Merianda et al., 2015).

### Axon growth analyses

For analyses of axon growth, dissociated DRGs grown on glass coverslips or glass bottom 24 well plates were immunostained with neurofilament antibodies as described above. Entire coverslips or bottoms of the 24 well plates were scanned with ImageXpress Micro system (Molecular Devices) using a 20X objective. Stitched images were assessed for total axon length, longest axon length, and branching using WIS-Neuromath (Rishal et al., 2013) or ImageJ with NeuronJ plugin programs (ImageJ plugin; NIH). All neurons where axonal arbors could be traced by the software were assessed (≥ 75 neurons per group/per experiment).

### Fluorescence recovery after photobleaching (FRAP)

FRAP analyses was performed with minor modifications (Vuppalanchi et al., 2010). DRG neurons were transfected with GFP^MYR^3’prenyl-Cdc42 or GFP^MYR^3’palm-Cdc42 or GFP^MYR^3’γ-actin as a negative control. Cells were maintained at 37°C, 5 % CO_2_ during imaging sequences. The 488 nm laser line on Leica SP8X confocal microscope was used to bleach GFP signal (Argon laser at 70 % power, pulsed every 0.82 sec for 80 frames). Pinhole was set to 3 Airy units to ensure full thickness bleaching and acquisition (63 X/1.4 NA oil immersion objective) (Yudin et al., 2008). Prior to photobleaching, neurons were imaged every 60 sec for 2 min to acquire baseline fluorescence in the region of interest (ROI; 15 % laser power, 498-530 nm for GFP). The same excitation and emission parameters were used to assess recovery over 15 min post-bleach with images acquired every 30 sec. To determine if fluorescence recovery in axons due to translation, DRG cultures were treated with 100 µM anisomycin (Sigma) for 30 min prior to photobleaching.

### Analysis of RNA-Seq data

Raw FASTQ files of GSE51572, GSE66230, and GSE67828 were downloaded from NCBI GEO database and adaptor sequences were trimmed as necessary (Briese et al., 2016; Minis et al., 2014; Taliaferro et al., 2016). Unpublished axonal RNA-seq data from cultures of adult C57Bl/6 mouse DRGs was similarly analyzed. Reads were then aligned to mm10 genome using STAR with default parameters. Mapped reads for mRNAs were counted by HTSeq and then normalized by trimmed mean of M values using EdgeR. Normalized counts were used to quantitate levels of *Cdc42* splice variants in axons, neurites or cell bodies.

### Experimental design, image analyses, and statistical analyses

All experiments using cultured neurons were designed to account for technical and inter-experimental variations. Consequently, all imaging experiments included at least three technical replicates on each culture (*i*.*e*., separate coverslips) and experiments were replicated across at least 3 separate culture preparations. High content imaging was used for neurite outgrowth analyses such that at least 75 neurons per coverslip were analyzed. For molecular studies of transfected cultures (*i*.*e*., for ddPCR), analyses were performed on at least three separate culture preparations. Positive and negative controls were included as outlined above and in the Results section.

All FISH/IF performed on tissue sections were imaged by confocal microscopy using a Leica SP8X confocal microscope with HyD detectors and post-processing measures to distinguish axonal signals from non-neuronal signals. Scrambled probes were used to assign image acquisition parameters to limit any nonspecific signal from the probes. A 145 x 145 μm nerve segment was scanned by taking xyz image stacks with 63x oil-immersion objective (1.4 NA). This xyz plane stack was captured at two locations along each nerve section. The *Colocalization Plug-in* for NIH *ImageJ* (https://imagej.nih.gov/ij/plugins/colocalization.html) was used to extract RNA signals from FISH probes in each optical plane overlapping with axonal markers (NF) in the xy plane (Terenzio et al., 2018). All FISH signal quantifications for axonal mRNA signals from tissue culture sections were generated by analyses of pixel intensity across each xy plane of the extracted ‘axon only’ channels for the image sequences using Image J. These FISH signal intensities across the individual xy planes were then normalized to the area of NF immunoreactivity in each XY plane and averaged across the image Z stack of the tile images (Kalinski et al., 2015). The relative mRNA signal intensity was averaged for all tiles in each biological replicate.

For quantification of pixel intensity distribution in growth cones and terminal axon shaft and growth cones were imaged by confocal microscopy using a Leica SP8X confocal microscope with HyD detectors and deconvolved using Huygens Deconvolution software (https://svi.nl/Huygens-Deconvolution). Imaging sequences for terminal axon shaft and growth cones consisted of acquiring a 40-50 μm segment by taking *xyz* image stacks with 63× oil-immersion objective (1.4 NA) for 23-27 optical planes at *z* depth of 5.8-6.8 μm (0.25 μm interval between planes). The image stacks were projected using the maximum XYZ projection. For unbiased protein distribution profiling across the growth cone, lines of 20 µm width were drawn using the line tool in Image J and tracked 15 µm proximally in to the axon shaft starting from the growth cone tip. Surface plots of the tracked growth cones were measured using ImageJ. For peripheral, central, and proximal growth cone protein quantification segmented line tool in ImageJ was used to manually track these areas and integrated pixel intensities were measured.

Fluorescent intensities in the bleached regions of interest (ROI) were calculated by the Leica LASX software. For normalizing across experiments, fluorescence intensity value at t = 0 min post-bleach from each image sequence was set as 0 %. The percentage of fluorescence recovery at each time point after photobleaching was calculated by normalizing relative to the pre-bleach fluorescence intensity for each image sequence was set at 100 %.

Quantitative data are reported as mean ± standard deviation (SD) or standard error of the mean (SEM) as indicated in the figure legends. Student’s t-test or one-way ANOVA with pairwise comparisons and Tukey post-hoc was used to determine significance differences between groups as indicated in the figure legends. GraphPad Prism 5 software was used for all statistical analyses.

## ACKNOWLWDGEMENTS and FUNDING

This work was supported by grants from NIH (R01-NS089663 to JLT; R01-CA111891 and R15-CA165202 to QL), the Dr. Miriam and Sheldon G. Adelson Medical Research Foundation (to JLT and GC), the Wings for Life - Spinal Cord Research Foundation (WFL-US-09/18 to PP), South Carolina Spinal Cord Injury Research Fund (2019 PD-02 to PKS), the Harriet and John Wooten Foundation for Alzheimer’s Disease Research (to QL), and ASPIRE award from the University of South Carolina Office of Research (to SJL and ANK). JLT is the incumbent SmartState Chair in Childhood Neurotherapeutics at the University of South Carolina.

## AUTHOR CONTRIBUTIONS

SJL, ANK, and JLT designed the study. SJL, ANK, MDZ, RK, PP, PKS, KDL and CRM performed the experiments and analyzed the results. QL and BJA provided critical reagents and advice. SJL, ANK, MDZ, PKS, and JLT wrote the article with inputs from PP, QL, RK and GC.

## COMPETING INTERESTS

The authors declare that they have no conflict of interest.

## SUPPLEMENTAL MATERIALS

**Supplemental Table S1:** Alternatively spliced variants of Cdc42 mRNA are found from fish to mammals. Sequence comparison for indicated species shows conservation of *Cdc42* alternative splicing from fish to primates.

| **Species (common name)** | **NCBI Accession #**<br>***prenyl-Cdc42*** | **NCBI Accession #**<br>***palm-Cdc42*** | **CaaX motif**<br>***prenyl-Cdc42*** | **CCaX motif**<br>***palm-Cdc42*** |
| --- | --- | --- | --- | --- |
| *Saccharomyces cerevisiae* (yeast) | NM_001182116.1 | - | CAIL | - |
| *Caenorhabditis elegans* (worm) | NM_063197.7 | - | CNIL | - |
| *Saccoglossus kowalevskii* (worm) | NM_001168041.1 | - | CVLL | - |
| *Acanthaster planci* (star fish) | XM_022244898.1 | - | CSLL | - |
| *Danio rerio* (zebra fish) | NM_200632.2 | NM_001018120.2 | CVLL | CCIF |
| *Xenopus tropicalis* (frog) | NM_001017070.3 | NM_001008026.1 | CRLL | CCIF |
| *Xenopus laevis* (frog) | NM_001085899.1 | XM_018224258.1 | CMLL | CCIF |
| *Nanorana parkeri* (frog) | XM_018568983.1 | XM_018568984.1 | CRLL | CCIF |
| *Callorhynchus milii* (shark) | XM_007897764.1 | XM_007897763.1 | CVLL | CCIF |
| *Python bivittatus* (snake) | XM_015889191.1 | XM_015889192.1 | CVLL | CCIF |
| *Alligator mississippiensis* (alligator) | XM_014597185.2 | XM_014597186.2 | CVLL | CCIF |
| *Gallus gallus* (chicken) | NM_205048.1 | XM_015296827.1 | CVLL | CCIF |
| *Chelonia mydas* (turtle) | XM_007065922.1 | XM_007065923.1 | CVLL | CCIF |
| *Mus musculus* (mouse) | NM_009861.3 | NM_001243769.1 | CVLL | CCIF |
| *Homo sapiens* (human) | NM_001039802.1 | NM_044472.2 | CVLL | CCIF |

**Supplemental Table S2:** Normalized counts for prenyl-Cdc42 and palm-Cdc42 mRNAs from subcellular transcriptome datasets of different neuron types. Raw data of published and unpublished subcellular transcriptome datasets were performed by RNA-seq analysis pipeline (STAR, HTSeq and EdgeR) to obtain the normalized counts. Datasets were from cell body and axon-enriched transcriptomes of: embryonic DRG neurons (Minis et al., 2014), embryonic motor neurons (Briese et al., 2016), adult DRG neurons (unpublished data), and cell body-/neurite-enriched transcriptome of embryonic cortical neurons (Taliaferro et al., 2016).

| Neuron (species) | Cell Body Compartment (FKPM ± SEM) | | Axon/Neurite Compartment (FKPM ± SEM) | |
| --- | --- | --- | --- | --- |
|  | *Palm-Cdc42* | *Prenyl-Cdc42* | *Palm-Cdc42* | *Prenyl-Cdc42* |
| Embryonic DRG (mouse) | 109.3 ± 23.9 | 271.2 ± 16.1 | 47.6 ± 2.3 | 384.7 ± 6.1 |
| Embryonic motor (mouse) | 105.2 ± 10.0 | 191.4 ± 14.4 | 18.0 ± 5.6 | 96.7 ± 40.0 |
| Adult DRG (mouse) | 13.0 ± 1.8 | 67.8 ± 6.6 | 4.7 ± 2.1 | 46.6 ± 19.3 |
| Embryonic cortical (mouse) | 428.1 ± 145.9 | 945.6 ± 7.3 | 343.6 ± 62.7 | 908.0 ± 44.8 |

## SUPPLEMENTAL FIGURE LEGENDS

**Supplemental Figure S1:**
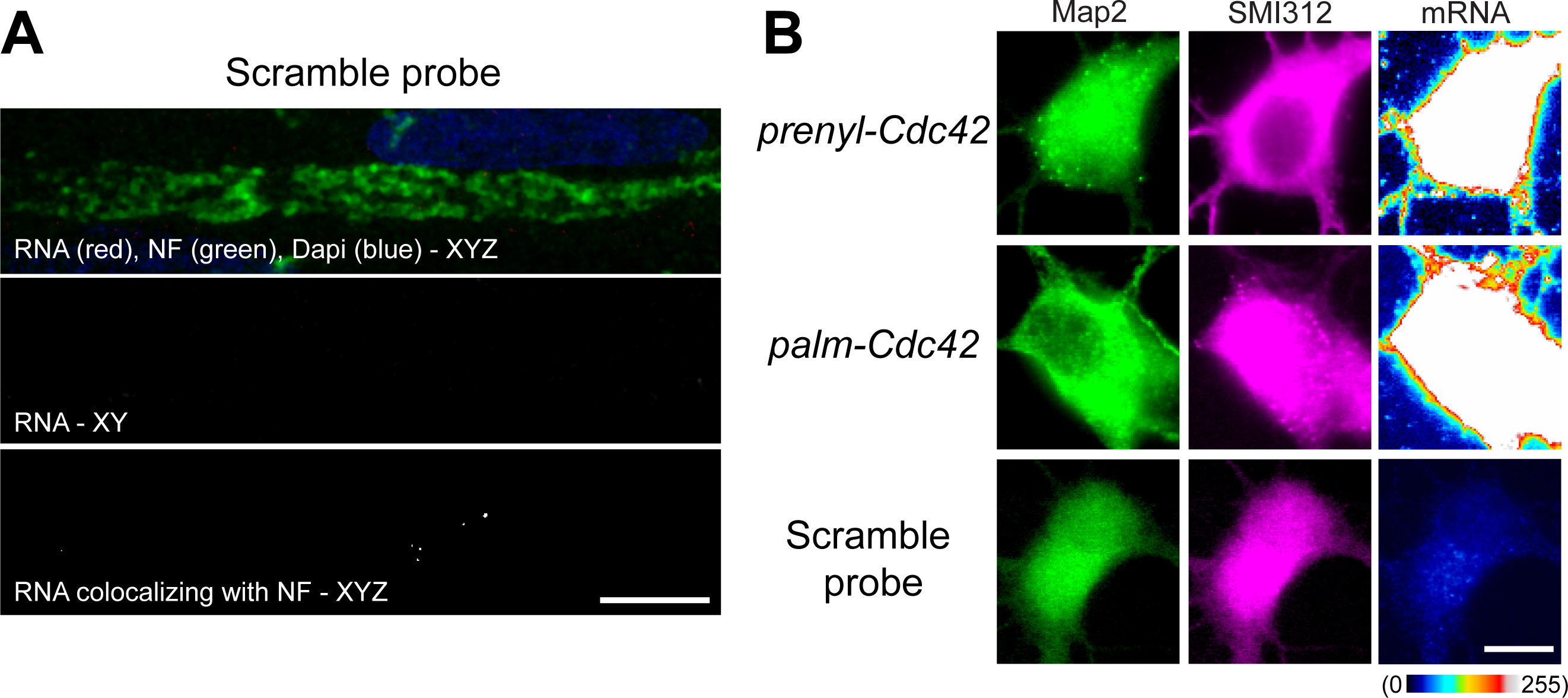
Prenyl-Cdc42 mRNA is localizes to axonal processes. **A**, Representative FISH/IF images for scrambled RNA FISH probe for sciatic nerve from experiment shown in Figure 2A [scale bar = 10 µm]. **B**, Representative FISH/IF images for cell bodies of cortical neurons corresponding to the axon and dendrite images from Figure 3D-E are shown [scale bar = 20 µm].

**Supplemental Figure S2:**
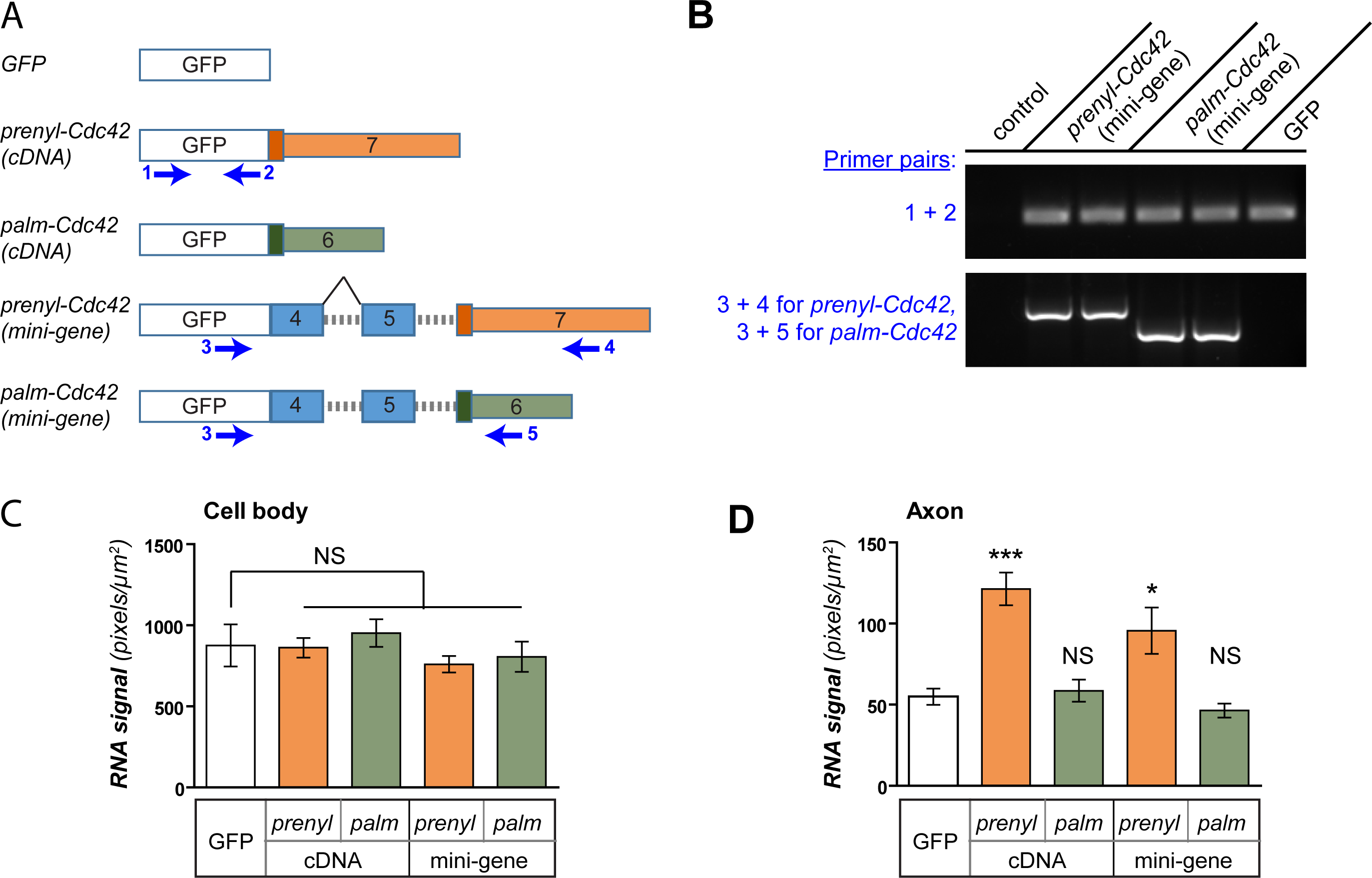
Axonal localization of prenyl-Cdc42 does not require splicing. **A**, Schematics for GFP-Cdc42 constructs designed to mimic pre-mRNA splicing differences between *prenyl-Cdc42* and *palm-Cdc42* mRNAs (‘mini-genes’) vs. mature mRNA (‘cDNA’). Sites for annealing of primers 1-5 (blue text) are shown below each RNA schematic. **B**, Representative ethidium-stained agarose gel for RT-PCR to detect splice variants using primers outlined in A as indicated. The two lanes for the *CDC42* mini-genes show replicate transfections. Amplification with primers 1 + 2 shows relatively equivalent expression across the transfections. Amplification with primers 3 + 4 and 3 + 5 shows anticipated splice variants for mini-gene constructs. Control is untransfected DRGs. **C-D**, Quantification of GFP mRNA signals from FISH for cell bodies (**C**) and axons (**D**) is shown as mean ± SEM. GFP was used as a negative control (N ≥ 30 neurons from 3 biological replicates; * p < 0.05, *** p < 0.001 by one-way ANOVA with pair-wise comparison and Tukey post-hoc).

**Supplemental Figure S3:**
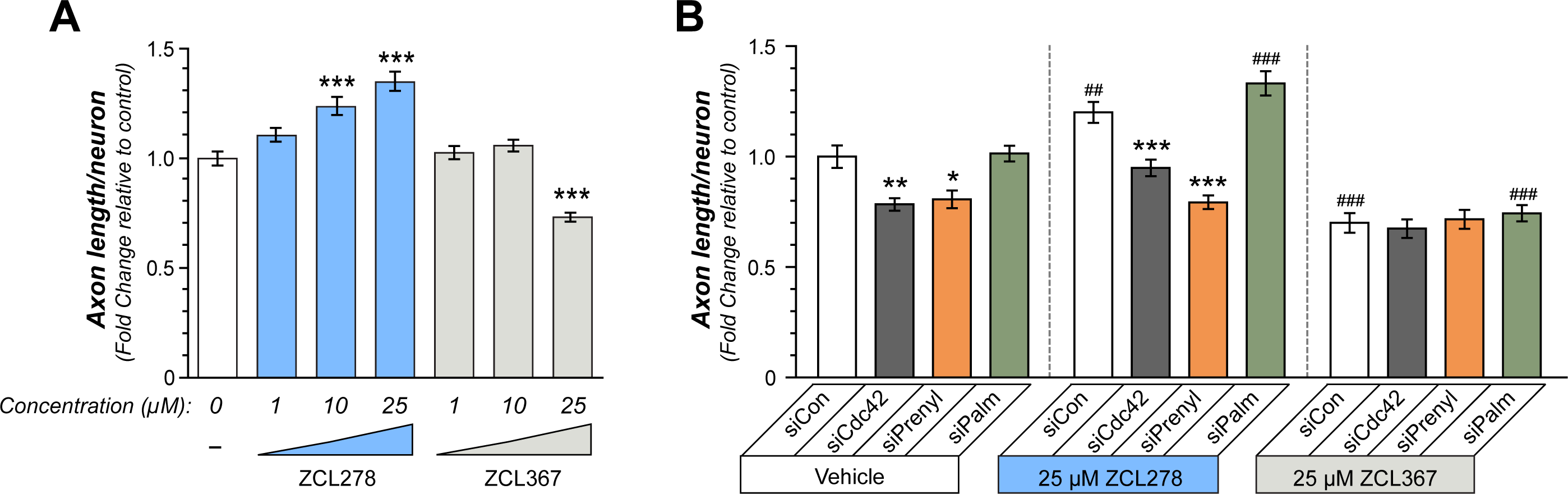
Axon growth enhancement by prenyl-Cdc42 requires Cdc42 activity. **A**, Total axon length per neuron for DRG cultures after 48 hours exposure to ZCL278 and ZCL367 shown as mean ± SEM relative to vehicle (control; N ≥ 200 neurons in 3 independent experiments; *** P < 0.005 by one-way ANOVA with pair-wise comparison and Tukey post-hoc tests). **B**, Total axon length for DRGs transfected with indicated siRNAs and treated with ZCL278 or ZCL367 is shown as mean ± SEM relative to siCon + vehicle (N ≥ 150 neurons in three independent experiment; * p < 0.05, ** p < 0.01, and *** p < 0.005 for comparison to siCon within vehicle, ZCL278, and ZCL367 groups and # p < 0.05, ## p < 0.01, ### p < 0.005 group-wise comparisons of ZCL278 or ZCL367 groups to vehicle group for each corresponding siRNAs by one-way ANOVA with pair-wise comparison and Tukey post-hoc tests; ZCL367 + pan-siCdc42, siPrenyl or siPalm are not significantly different than ZCL367 plus siCon).

## Notes

### Competing Interest Statement

The authors have declared no competing interest.

### Summary of Updates

Figure 2 - Addition of in situ hybridization data for Cdc42 mRNAs in vivo. Figure 6 - Quantification of CDC42-GFP protein levels in growth cones.

